# New graph-theoretical-multimodal approach using temporal and structural correlations reveal disruption in the thalamo-cortical network in patients with schizophrenia

**DOI:** 10.1101/489997

**Authors:** Paolo Finotelli, Caroline Garcia Forlim, Paolo Dulio, Leonie Klock, Alessia Pini, Johanna Bächle, Laura Stoll, Patrick Giemsa, Marie Fuchs, Nikola Schoofs, Christiane Montag, Jürgen Gallinat, Simone Kühn

**Affiliations:** Politecnico di Milano, Department of Mathematics “F. Brioschi”, Piazza L. da Vinci 32, 20133, Milan, Italy; University Medical Center Hamburg - Eppendorf, Clinic and Policlinic for Psychiatry and Psychotherapy, Martinistraße 52, 20246, Hamburg, Germany; Università Cattolica del Sacro Cuore, Department of Statistical Sciences, Largo A. Gemelli, 1 – 20123 Milan, Italy; Charité University Medicine and St. Hedwig-Krankenhaus, Department of Psychiatry and Psychotherapy, Große Hamburger Straße 5-11, 10115 Berlin, Germany; Max Planck Institute for Human Development, Center for Lifespan Psychology, Lentzeallee 94, 14195 Berlin, Germany

**Keywords:** network analysis, functional connectivity, structural connectivity, resting state fMRI, VBM, schizophrenia, thalamo-cortical network, dysconnectivity

## Abstract

Schizophrenia has been understood as a network disease with altered functional and structural connectivity in multiple brain networks compatible to the extremely broad spectrum of psychopathological, cognitive and behavioral symptoms in this disorder.

When building brain networks, functional and structural networks are typically modelled independently: functional network models are based on temporal correlations among brain regions, whereas structural network models are based on anatomical characteristics. Combining both features may give rise to more realistic and reliable models of brain networks.

In this study, we applied a new flexible graph-theoretical-multimodal model called FD (F, the functional connectivity matrix, and D, the structural matrix) to construct brain networks combining functional, structural and topological information of MRI measurements (structural and resting state imaging) to patients with schizophrenia (N=35) and matched healthy individuals (N=41). As a reference condition, the traditional pure functional connectivity (pFC) analysis was carried out.

By using the FD model, we found disrupted connectivity in the thalamo-cortical network in schizophrenic patients, whereas the pFC model failed to extract group differences after multiple comparison correction. We interpret this observation as evidence that the FD model is superior to conventional connectivity analysis, by stressing relevant features of the whole brain connectivity including functional, structural and topological signatures. The FD model can be used in future research to model subtle alterations of functional and structural connectivity resulting in pronounced clinical syndromes and major psychiatric disorders. Lastly, FD is not limited to the analysis of resting state fMRI, and can be applied to EEG, MEG etc.

## Introduction

Schizophrenia is a mental illness with heterogeneous symptoms including positive symptoms such as delusions and hallucinations and negative symptoms such as reduced emotional expression, lack of motivation, among others. It has been hypothesised that schizophrenia could be understood as a network disease with dysfunctional connectivity between multiple brain regions (Friston and Frith 1995; Andreasen et al. 1998; Friston 1999). According to this hypothesis, symptoms are assumed to emerge from a failure in the functional integration of information processing in the brain (Garrity et al. 2007). Several studies have found abnormal functional and structural connectivity in multiple brain networks supporting the theory of schizophrenia as a dysconnectivity syndrome (Friston and Frith 1995; Hadley et al. 2016) and suggesting functional and structural disturbances in wide whole-brain-range connectivity (Nelson et al. 1998; Andreasen and Pierson 2008; Kühn et al. 2012; Zalesky et al. 2012; Cocchi et al. 2014; van den Heuvel and Fornito 2014; Barch 2014; Singh et al. 2015).

Brain networks in neuroimaging using functional magnetic resonance imaging (fMRI) are composed by brain areas, called nodes, and the connections between nodes are called links (van den Heuvel and Hulshoff Pol 2010). Links in the brain networks can be constructed using structural or functional information. In structural networks, links are built using correlations of morphometric features from diffusion tensor imaging (DTI) and MRI as fibres in the white matter/total number of interconnecting streamlines, cortical thickness or grey matter volume. Whereas in functional networks, two regions are said to be functionally connected, and therefore, engaged in information processing, if they present a temporal correlation. Links constructed using these temporal correlations are usually calculated using Pearson’s correlation coefficient.

Typically, structural and functional brain networks are calculated and analysed separately, for example, functional networks are built using solely temporal correlations given by Pearson correlation function, regardless of any structural information. However, combining these two equally important features of brain function and morphology might give rise to more realist, complete and even reliable models of brain networks (Bowman et al. 2012; Xue et al. 2015; Calhoun and Sui 2016; Battiston et al. 2017; Kang et al. 2017; Calamante et al. 2017; Chu et al. 2018).

In this framework, a new graph-theoretical-multimodal model was proposed to study connectivity, that is called the FD model (F, the functional connectivity matrix, and D, the structural matrix (Finotelli and Dulio 2015). The FD model adds, to the pure functional connectivity (pFC), structural and topological information that can be obtained from multiple different techniques: topological information given by nodal degree, functional connectivity matrix given by Pearson’s correlation, Spearman correlation, mutual information, Granger causality or any other correlation/causality index, and for the structural matrix D, from morphometric characteristics, such as volume, cortical thickness and grey/white matter connectivity as well the Euclidean distances between pairs of brain areas, DTI data, etc. The FD model has been successfully applied to EEG data with subjects performing a musical task (Finotelli et al. 2016), and to synthetic data simulation human’s brain functional activity at rest (Dulio et al. 2018).

The aim of this paper is to analyze whole brain network of patients with schizophrenia and a matched group of healthy individuals by means of two different models. First, pFC model, commonly used in the literature, that takes only the functional connectivity into consideration. Second, the FD model that utilizes not only the functional data but also structural and topological properties of the cerebral network.

## Methods

### Participants

Forty-one were healthy individuals and thirty-five individuals who met the criteria for a diagnosis of schizophrenia following the International Classification for Diseases and Related Health Problems (ICD-10) (World Health Organization 2012) were included in the study. The recruitment of patients diagnosed with schizophrenia took place at St. Hedwig Hospital, Department for Psychiatry and Psychotherapy of the Charité-Universitätsmedizin Berlin (Germany). A trained clinician assessed the severity of symptoms with the Scale for Assessment of Negative Symptoms (SANS) (Andreasen 1989), and Scale for Assessment of Positive Symptoms (SAPS) (Andreasen 1984). For the healthy individuals, the recruitment was accomplished using advertisements and flyers. Healthy individuals did not meet the criteria for any psychiatric disorder based on information acquired with the Mini International Neuropsychiatric Interview (MINI) (Ackenheil et al. 1999) and were not in current or past psychotherapy of an ongoing mental health-related problem. Healthy individuals matched the group of patients in terms of age, sex, handedness and level of education (Table 1). Handedness was acquired using the Edinburgh Handedness Inventory, cognitive functioning was tested using the Brief Assessment of Cognition in Schizophrenia (Keefe et al., 2008) and verbal intelligence with a German Vocabulary Test (Schmidt & Metzler, 1992). All procedures of the study were approved by the ethics committee of the Charité-Universitätsmedizin Berlin.

**Table 1.** Demographics and clinical characteristics.

|  | **HC (n=41)**<br>Mean(Standard<br>Deviation) | **SCZ(n=35)**<br>Mean(Standard<br>Deviation) | *p*Value |
| --- | --- | --- | --- |
| Gender (male/female) | 24/17 | 21 /14 |  |
| Age (years) | 35.2 (11.0) | 35.3 (10.8) | 0.953 |
| Framewise displacement | 0.15(0.02) | 0.17(0.02) | 0.42 |
| Edinburg Handedness Inventory<sup>a</sup> | 79.4 (38.5) | 75.56 (52.6) | 0.742 |
| Education (years) | 14.1 (2.9) | 13.1 (3.8) | 0.183 |
| BACS<sup>a</sup> | 270.4 (37.0) | 234.1 (28.4) | < 0.001 |
| Verbal Intelligence (IQ)<sup>a</sup> | 100.5 (10.6) | 94.4 (12.8) | 0.031 |
| Illness duration (years) |  | 9.4 (8.8) |  |
| Illness onset (age in years) |  | 25.6 (8.9) |  |
| Chlorpromazine-equivalent (mg) |  | 317.1 (221.6) |  |
| SANS Composite Score<sup>a</sup> |  | 20.2 (12.0) |  |
| SAPS Composite Score<sup>a</sup> |  | 15.4 (14.9) |  |
| BACS, Brief Assessment of Cognition in Schizophrenia; SAPS, Scale for Assessment of Positive Symptoms; SANS, Scale for Assessment of Negative Symptoms;<br><sup>a</sup> sum score of items reported. |  |  |  |

### MRI Data acquisition

Images were collected on a Siemens Tim Trio 3T scanner (Erlangen, Germany) with a 12-channel head coil. Structural images were obtained using a T1-weighted magnetization prepared gradient-echo sequence (MPRAGE) based on the ADNI protocol (TR=2500ms; TE=4.77ms; TI=1100ms, acquisition matrix=256×256×176; flip angle = 7°; 1x1x1mm^3^ voxel size). Whole brain functional resting state images during 5 minutes were collected using a T2*-weighted EPI sequence sensitive to BOLD contrast (TR=2000ms, TE=30ms, image matrix=64×64, FOV=216mm, flip angle=80°, slice thickness=3.0mm, distance factor=20%, voxel size 3×3×3mm^3^, 36 axial slices). Before resting state data acquisition was started, participants were in the scanner for about 10 minutes during which a localizer and the anatomical images were acquired so that subjects could get used to the scanner noise. During resting state data acquisition participants were asked to close their eyes and relax during data acquisition.

### Preprocessing of resting state data

The first 5 images were discarded to ensure for steady-state longitudinal magnetization. The data was then corrected for slice timing and realigned. Individual T1 images were coregistered to functional images and segmented into gray matter, white matter, and cerebrospinal fluid. Data was spatially normalized to the MNI template and, to improve signal-to-noise ratio, spatially smoothed with a 6-mm FWHM. Motion and signals from white matter and cerebrospinal fluid were regressed. Data was then filtered (0.01 – 1 Hz) to reduce physiological high-frequency respiratory and cardiac noise and low-frequency drift and, finally, detrended. All steps of data preprocessing were done using SPM12 except filtering that was applied using the REST toolbox (Song et al. 2011). In addition, to control for motion, the voxel-specific mean framewise displacement was calculated ( Power and colleagues (Power et al. 2012). Framewise displacement values were below the default threshold of 0.5 for control and patient group (table 1).

### Voxel-based morphometry (VBM)

VBM, an unbiased objective technique, has been developed to investigate regional differences in the brain anatomy (Mechelli et al. 2005) using MRI. VBM estimates regional white and/or gray matter volume. Structural data was processed by means of the VBM8 toolbox (http://dbm.neuro.uni-jena.de/vbm.html) and SPM8 (http://www.fil.ion.ucl.ac.uk/spm) with default parameters. The VBM8 toolbox involves bias correction, tissue classification and affine registration. The affine registered gray matter (GM) and white matter (WM) segmentations were used to build a customized DARTEL diffeomorphic anatomical registration through exponentiated lie algebra template. Then warped GM and WM segments were created. Modulation was applied in order to preserve the volume of a particular tissue within a voxel by multiplying voxel values in the segmented images by the Jacobian determinants derived from the spatial normalization step. In effect, the analysis of modulated data tests for regional differences in the absolute amount (volume) of GM.

### The mathematical models

In this section we introduce the two models involved in this study: pFC and FD, and describe the basic concepts of Graph theory and Linear Algebra (the matrices).

#### Network representation

Adjacency matrix is the mathematical expression of a network. In brain networks, each row and column is a neural element and how they are related are the weights. If the weights are obtained calculating the pFC between n neural elements, e.g. voxels or brain regions, the resulting adjacency matrix will have *n* rows and *n* columns, being called a square matrix. Importantly, by exchanging the rows with the columns in a matrix **A** returns a matrix **A^T^** called the transpose of **A**. **A^T^** is so characterized by having *n* rows and *n* columns. If **A**=**A^T^** the matrix is said to be symmetric, which holds to pFC but not to effective connectivity due to the different nature of the measure.

#### Degree of a node

The degree of a neural element (or node) is the sum of all its connections in the network. This topological metric is useful to get information about the centrality of the node in the network and a first step in understanding if the node can take on the role of a hub. Hubs play a central role in the network, integrating and distributing information in very effective ways due to the number and positioning of their connections in a network.

Formally, a graph *G* refers to a set of vertices (or nodes) and of edges (or links) that connect the vertices. Two nodes are said to be adjacent if they are endpoints of an edge. An important number associated with each vertex is its degree. The degree of an arbitrary node *v* is the number *deg(v)* of the adjacent nodes. In our study, the nodes represent the cerebral areas in the AAL template. Importantly, a graph *G* can be equivalently represented by a square matrix **AG**. This is done by defining a weight for each edge of the graph *G*. For each *i* ∈{1,2,…,m} and *j* ∈{1,2,…,n} the entry (i,j) of the matrix **A_G_** is the weight of the edge connecting the nodes *i* and *j*, if it exists, or 0 otherwise.

The degree of the nodes can be calculated by adding all the non-zero elements of the columns (or equivalently, due to the symmetry, of the rows) of the connectivity matrix associated to that node.

#### The pFC model

Functional connectivity computed based on Pearson correlation coefficients is the most frequently applied method in the literature. This is a measure of linear correlation between two variables X and Y. The correlation values range from -1 to +1. When applied to neuroimaging, the correlations refer to a statistical dependence between physiological recordings that have been acquired from distinct neural elements (nodes). In fMRI, the neural elements are most likely brain regions or voxels and the physiological recordings are the indirect measure of the neural activity: the blood-oxygen-level dependent (BOLD). The Pearson correlation coefficient is symmetric, that is, correlation between X and Y is the same as the correlation between Y and X: corr(X,Y)=corr(Y,X). This leads to an important property for the mathematical description of a network: the adjacency matrix is symmetric.

#### The FD model

The FD model combines functional information given by the pFC matrix with structural and topological information: the FD model computes the connectivity as a function of, not only the strength of the statistical correlations as in pFC, but also as a function of the node degrees, and the structural connectivity (for details, see (Finotelli and Dulio 2015).

In the FD model, the general formula for calculating the functional weight of an arbitrary entry *W*_*i,j*_*(t)* of the matrix **W** is:

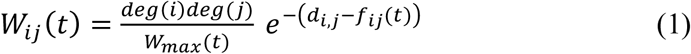

Where deg is the degree calculated as explained above, di,j and fij are, respectively, the generic entry of position (i,j) in the matrix **D**, which incorporates the structural information of the brain network, and in the matrix **F**, the functional connectivity matrix, whose entries represent the statistical dependence between the time series of measured neurophysiological signals and t is the time. Wij is within the range of 0 and +1, and therefore comparable to pFC. The exponent is the difference of two matrices, **D** and **F**, that should be characterized by having their entries in (nearly) the same range of variability. The FD model can employ both structural and functional data obtained from multiple different techniques. For more details concerning the explicit structure of (1) we refer the reader to (Finotelli and Dulio 2015). The only precaution is that **D** should be normalized in order to be comparable to matrix **F**.

In the present study we constructed the functional connectivity matrices **F** using Pearson’s correlation, the most popular connectivity measure, and the structural connectivity matrix **D** using VBM estimates of regional gray matter volume, resulting in a structural connectivity pattern matrix (He et al. 2007). VBM is a function of cortical thickness and cortical area, and therefore an important unbiased morphometry measure.

### Connectivity Matrices

#### Functional connectivity matrix generation

for each subject, the time series of each brain area (node) segmented according to the AAL 90 atlas (Tzourio-Mazoyer et al. 2002), were extracted using REST toolbox (Song et al. 2011). Then, the linear correlation between all pair of nodes (i,j=1,…,90) was calculated using Pearson’s correlation coefficient, resulting in a matrix **F**=[f_i,j_]. Two nodes were considered connected, and therefore a link set to fi,j, if the Pearson’s correlation coefficient was statistically significant. Due to the FD model, negative and self-correlations were not considered: negative f_i,j_ was set to 0 as well as all the elements on the principal diagonal of the matrices. Please note that the pFC is composed only by the **F** matrices, whereas the FD model follows formula (1).

#### Structural connectivity matrix generation

VBM was calculated as explained above. As for functional connectivity matrix, regions were segmented into the same 90 AAL regions as mentioned above. Two regions (nodes i,j=1,…,90) were assumed to be connected if the correlation of regional grey matter across subjects d_ij_ given by Pearson’s correlation coefficient was statistically significant, resulting in a structural connectivity pattern matrix **D**=[d_i,j_] for each group (He et al. 2007). Likewise, we did not consider negative and self-correlations due to the FD model: negative d_ij_ was set to 0 as well as all the elements on the principal diagonal of the matrices.

#### The FD model: Matrix generation

for each subject, the entries wi,j (i,j=1,…,90) of the FD model matrices **W** were calculated using (1), where deg(i) and deg(j) are the degree of nodes i and j respectively, **D**=[d_i,j_], is the stuctural matrix, **F(t)**=[f_i,j_(t=1)]=[f_i,j_], is the (thresholded) FC matrix **F(t=1)=F** see (Finotelli et al. 2016). For every matrix **W**, the thresholding procedure is performed by dividing all the entries of **W**, W_ij_, by W_max_, the maximum value of **W** (for details see (Finotelli and Dulio 2015).

## Statistical analysis

In both pFC and FD model, we performed a permutation-based unpaired t-test in order to compare the control and the patient group. We tested the null hypothesis of equality between the distributions of patients and controls on link î-j against the alternative of difference in distribution between the two groups on the same link. The test was based on the computation of the squared t-test statistic over all possible permutations of the data with respect to units (regardless of the groups). The p-value of the test was computed as the proportion of permutations leading to a value of the test statistic higher or equal with respect to the one observed with the original data. The significance level of the test α=0.05. In total, we compared, since the matrices were symmetric, 4050 links (we recall that we have zero entries on the main diagonal since there is not self-looping, so we excluded the diagonal from the comparison).

Finally, to control for multiple comparison, we performed the Bonferroni-Holm correction at α=0.05. The Bonferroni-Holm correction controls the family-wise error rate over the family of all links. The statistical methodology was the same for both models

## Results

We calculated whole brain connectivity using two different methods, first pFC and second the new graph-theoretical-multimodal FD model that takes structural as well as topological information of the brain network into account. First, we presented the results using the pFC and second the results of the FD model followed by a comparison between models.

### pFC

Brain network using pFC was calculated for each subject (see Method section) in both control and patient group (average whole brain network per group in Fig. 1). Then, a pair-wise group comparison was performed, and the p values were thresholded at 5% and corrected for multiple comparison (Bonferroni). No statistically significant connections were found between healthy controls and schizophrenia patients (see below for further discussion on statistically significant links, respective thresholds and multiple comparison methods between pFC and FD model).

**Figure 1:**
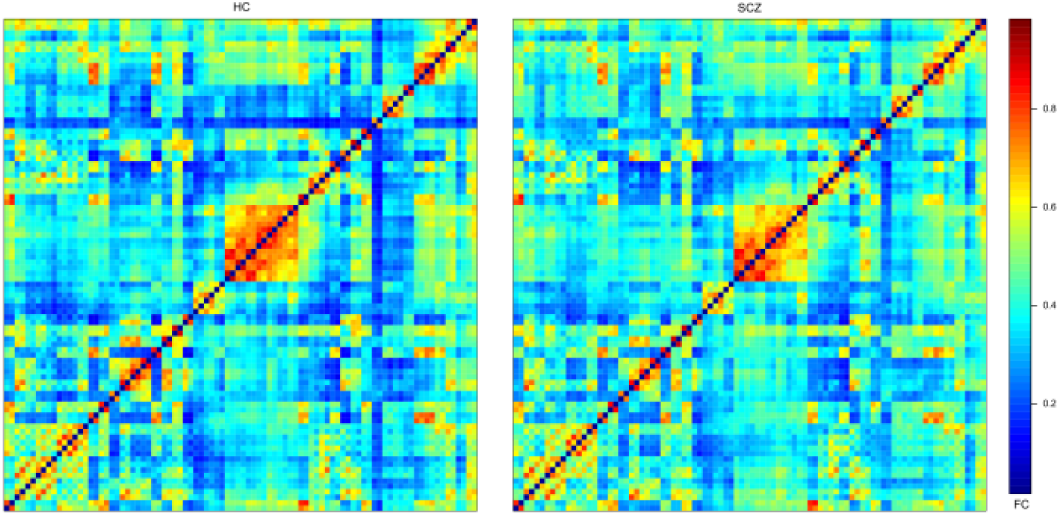
Average connectivity matrices calculated using pFC. The axis are the labels from the 90AAL template.

### FD model

A brain network was calculated for each subject (see Method section formula (1)). The average whole brain network per group is depicted in Fig. 2 - top. Then, pair-wise group comparisons were performed in the whole brain network, the p values (Fig. 2 - bottom) were thresholded at 5% and corrected for multiple comparison (Bonferroni).

**Figure 2:**
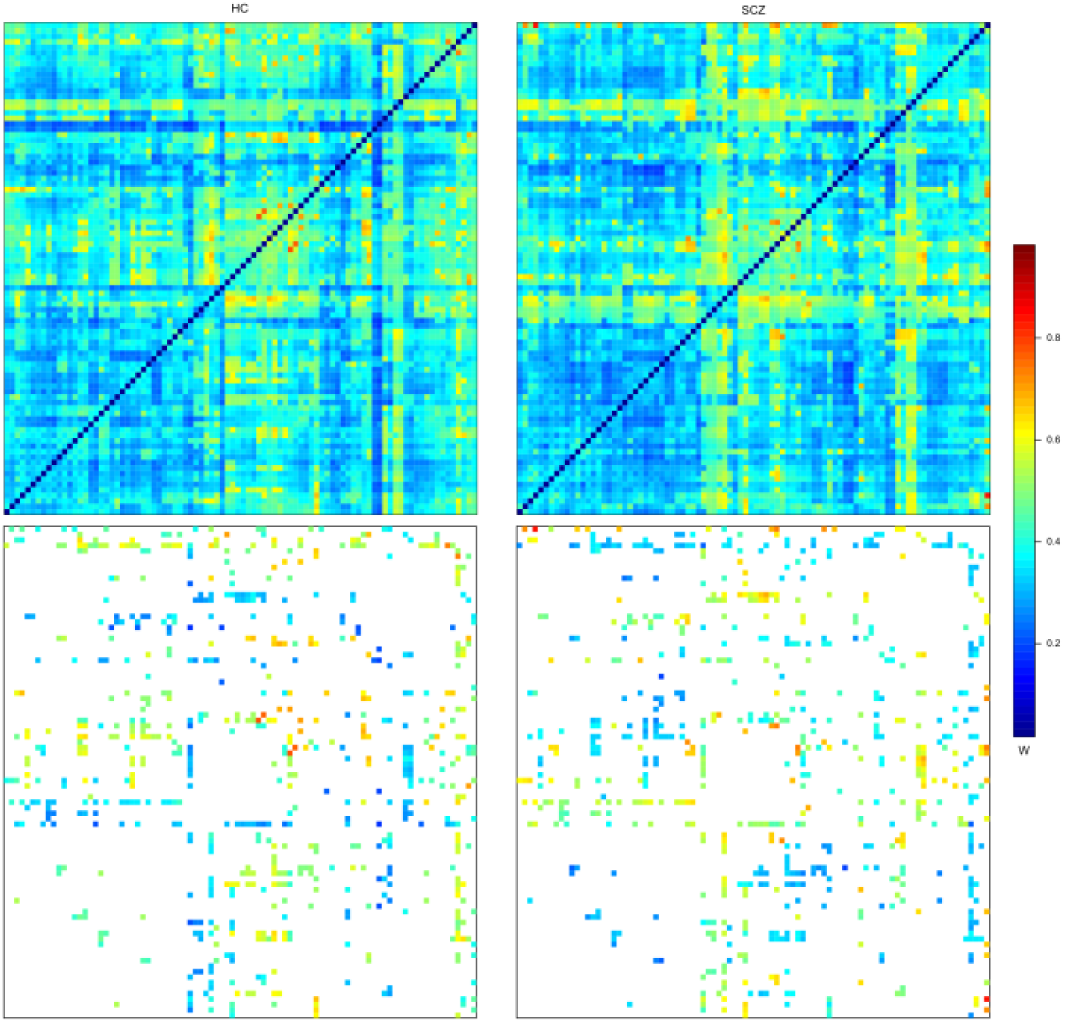
Top – Averaged connectivity matrices calculated using the FD model. Bottom – significant links after multiple comparison correction. The axis are the labels from the 90AAL template.

Connections that were statistically different between healthy control and schizophrenia patients, thresholded using the Bonferroni-adjusted p-value 0.05.

To explore the remaining links after the correction for multiple comparison, we reported the main links detected for the nodes with the highest connections (high degree). These nodes play a central role in the network given their interconnections and high degree (Fig. 3 and table 2). The brain regions that showed the most disrupted connections (high degree links) to other brain areas were the thalamus, inferior temporal gyrus, middle occipital gyrus, parahippocampus and superior occipital gyrus. The thalamus was the main hub connecting both cortical (mainly occipital and temporal via superior occipital gyrus) and subcortical areas (parahippocampal gyrus, hippocampus, precuneus). Impaired connections in frontal areas and sensorimotor cortex were also found. These links were mainly from/to inferior temporal and middle occipital. Interestingly, the hyperconnected subnetwork is part of a large network, known as thalamocortical network (Woodward et al. 2012).

**Table 2:** Impaired links established by the central nodes of the subnetwork.

| **Central nodes with impaired connectivity** | **Connected to:** |
| --- | --- |
| Right thalamus | Right hippocampus,<br>right parahippocampal gyrus,<br>bilateral calcarine,<br>bilateral cuneus,<br>bilateral lingual gyri,<br>right superior occipital gyrus, |
|  | left precuneus |
| Right inferior temporal gyrus | Right precentral,<br>right frontal superior,<br>right inferior frontal pars opercularis,<br>right frontal inferior pars triangularis,<br>right superior motor areas,<br>bilateral superior occipital gyri,<br>left inferial parietal gyrus,<br>right putamen |
| Right middle occipital gyrus | Right inferior frontal pars orbitalis,<br>left medial frontal pars orbitalis,<br>bilateral rectus,<br>bilateral paracentral lobuli,<br>left superior temporal pole |
| Right parahippocampus | Bilateral medial frontal pars orbitalis,<br>left anterior cingulum,<br>bilateral thalamus,<br>left middle temporal pole |
| Right superior occipital gyrus | Right inferior occipital gyrus,<br>right superior parietal gyrus,<br>bilateral thalamus,<br>bilateral inferior temporal gyrus |
| Left thalamus | Bilateral parahippocampal gyri,<br>left lingual gyrus,<br>right superior occipital gyrus,<br>right inferior occipital gyrus |

**Figure 3:**
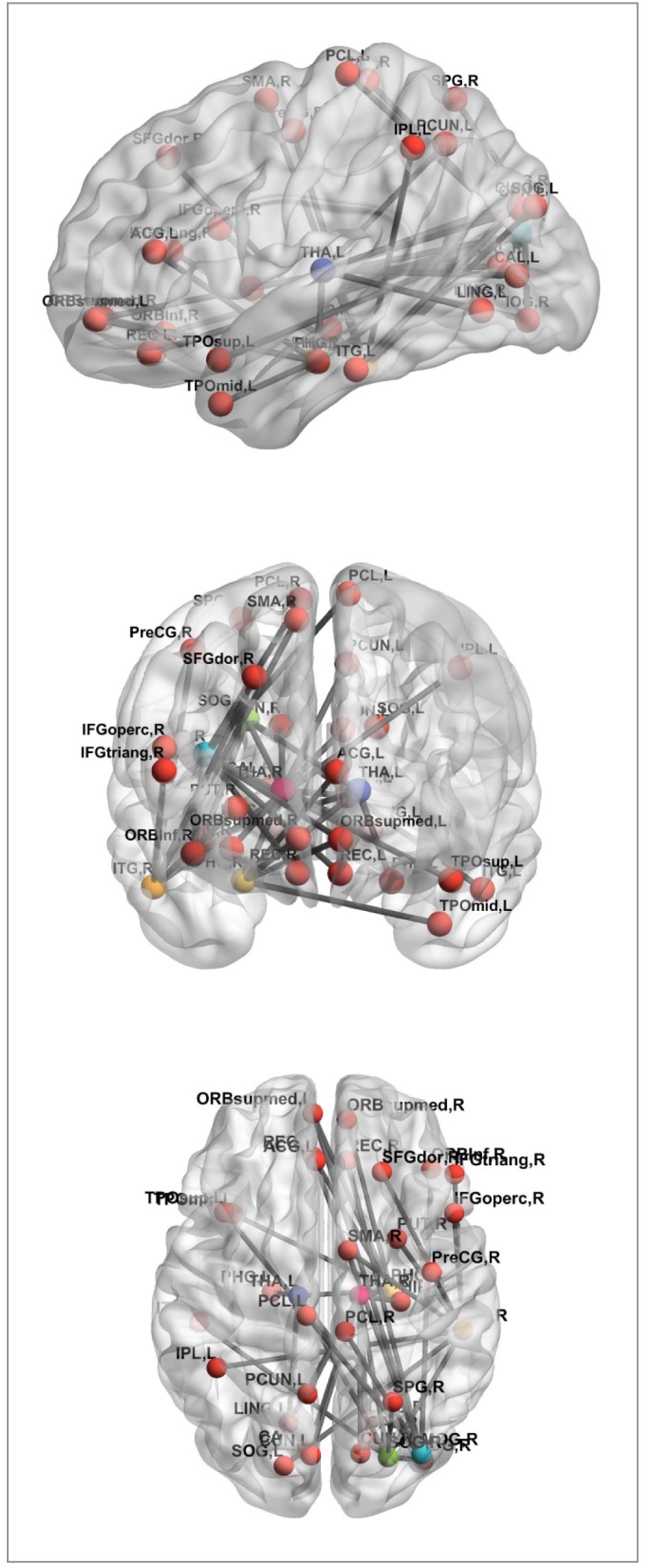
Impaired connections between healthy controls and schizophrenic patients. The central nodes of disrupted subnetworks were thalamus, inferior temporal gyrus, middle occipital gyrus, parahippocampal gyrus, superior occipital gyrus. The thalamus is considered the main hub of these central nodes connecting both cortical (mainly occipital and temporal via superior occipital gyrus) and subcortical areas (parahippocampal gyrus, hippocampus, precuneus). Impaired connections in frontal areas and sensorimotor cortex were mainly from/to inferior temporal and middle occipital.

(Figure 3 at the end of the paper)

### Model comparison: pFC and FD

We performed an exploratory analysis to compare the pFC and FD model, we showed the p-values of each connection resulting from the pairwise multiple comparison (Fig. 4 A). These matrices were thresholded as described above at 5% (Fig. 4B).

**Figure. 4:**
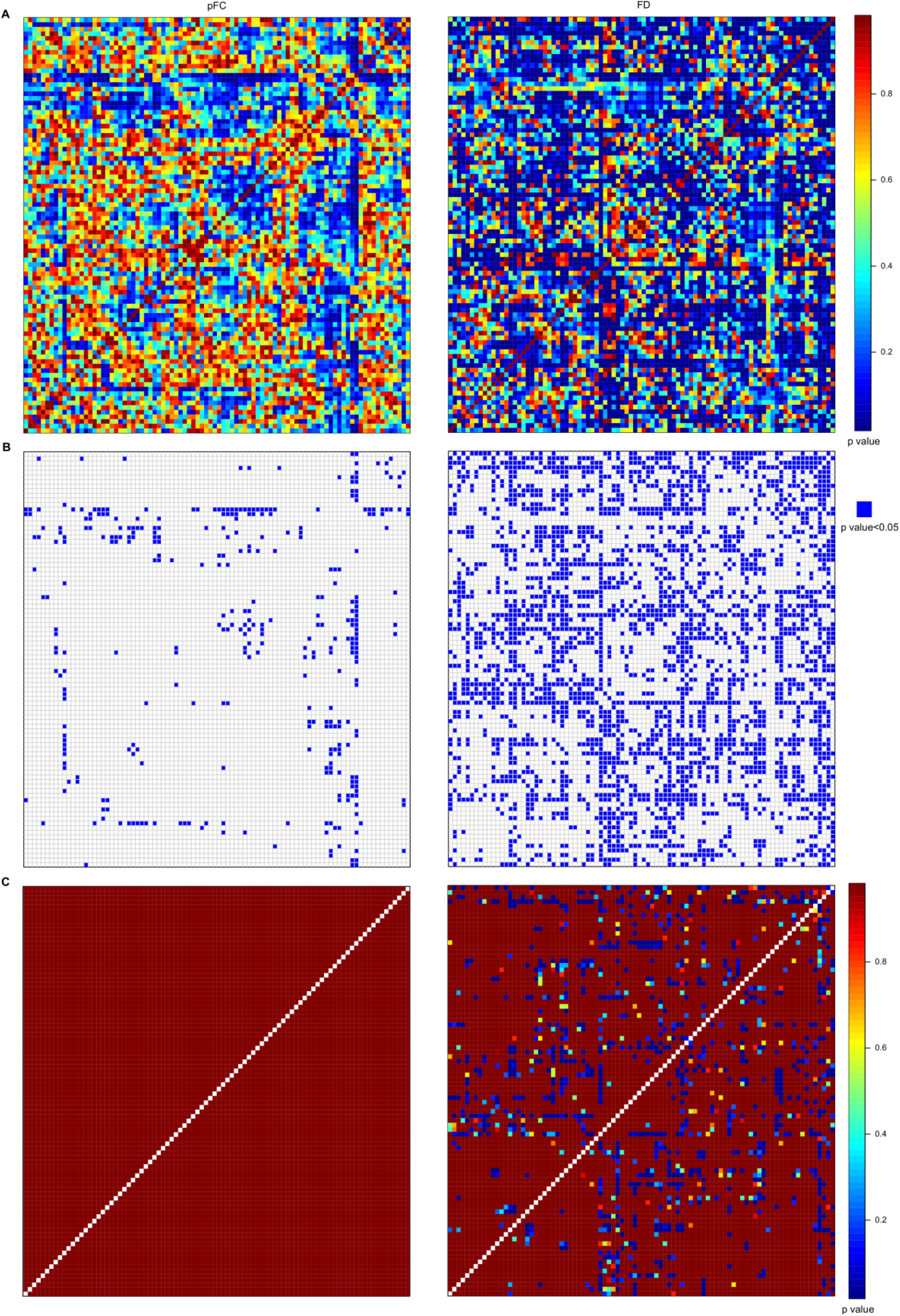
**A.** Adjacency matrix with uncorrected p values for FD model (right column) and pFC model (left column). **B.** Adjacency matrix with thresholded uncorrected p values at 5%. **C.** Bonferroni-Holm adjusted p-values for the FD model (right column) and the pFC model (left column). There were no significant differences in connectivity for the pFC model between healthy controls and schizophrenic patients. The axis are the labels from the 90AAL template.

In order to test whether the links found by applying the FD model were the same as the ones revealed by applying the standard pFC analysis, that is, whether there was a significant overlap between the links found by the two methods, we calculated the confusion matrix (Table 3).

**Table 3.** Confusion matrix – overlapping links between FD model and pFC model.

|  | Non Significant pFC | Significant pFC |
| --- | --- | --- |
| Non Significant W FD | **5099 (63%)** | 127 (1.6%) |
| Significant W FD | 2651 (33%) | **223 (2.8%)** |

Most of the links were non-significant for both models: 5099 (63%). The number of links that were significant for the FD model and for the pFC model was 223 (2.8%). The links that were significant for the FD model but not for the pFC model summed up to 2651 (33%). Significant links for pFC that were not in FD corresponded to 127 (1.6 %).

Finally, to adjust for the multiplicity of tests, we applied the Bonferroni-Holm (Fig. 4C) correction to the unadjusted p-values. After the adjustment, all p-values computed for the pFC model are equal to one. This means that using such a model no statistically significant differences can be detected between healthy controls and schizophrenia patients. However, when looking at results for FD model, there were statistically significant links after adjusting for multiple comparisons.

## Discussion

We applied a new model, called The FD model, to infer connectivity in the brain and compared it to the classical pFC. The main advantage of the FD model is that it takes into consideration not only the temporal correlations as in pFC does, but also includes information about structural connections and topological information of the network. Combining structural and functional information, two equally important features of brain, might give rise to more realistic and even reliable models of brain networks (Bowman et al. 2012; Xue et al. 2015; Calhoun and Sui 2016; Battiston et al. 2017; Kang et al. 2017; Calamante et al. 2017; Chu et al. 2018). In addition, degree is an important topological measure that reflects how well a specific region is connected to the whole network and it is therefore related to the notion of a hub. Hubs play a central role in the network, integrating and distributing information in effective ways due to the number and positioning of their connections in a network.

Once the models were applied to each subject in the schizophrenia group and in the healthy controls, individual whole brain networks were inferred followed by a pairwise comparison of all connections in the network between groups. In the FD model, we found wide-spread disrupted connectivity in key areas for schizophrenia (Andreasen 1997; Andreasen et al. 1998; Pergola et al. 2015), namely, hyperconnectivity in the thalamo-cortical network: thalamus, occipital, temporal parahippocampal gyrus and frontal areas. In this network there were six main central nodes: bilateral thalamus, right parahippocampal gyrus, right superior and middle occipital, right inferior temporal gyrus.

In addition, frontal and somatosensory areas were also present, in this cases the impaired connections arise mainly from the temporal inferior hub and middle occipital hub.

In line with previous models such as the “cognitive dysmetria” (Andreasen et al. 1998) and “filtering” models (Pergola et al. 2015), we observed that the thalamus was the main hub of the disrupted network. This region is located in a crucial anatomic position of the brain and its main function is the facilitation of connections as it receives input and output from distinct cortical areas. For this reason, it has been considered a key player in filtering and gating of information (Andreasen 1997; Pergola et al. 2015). Furthermore, it has been suggested that the thalamus might play a crucial role in schizophrenia because any disconnection in such a gate could lead to major changes in information load (Andreasen 1997) which in turn might account for the great variety of cognitive and clinical characteristic of the disease (Andreasen et al. 1998). Motivated by patients’ description of “*being bombed by stimuli which they have difficulty screening out”* (McGhie and Chapman 1961), Andreasen (Andreasen 1997) hypothesed a scenario in which information overload due to filtering and gating dysfunction could lead to a misinterpretation of the “self/not self”, auditory hallucinations, persecution, lack of energy etc, that represent key symptoms of schizophrenia. Although a growing body of evidence points to disruptions in the thalamus in schizophrenia patients, (Pergola et al. 2015; Ferri et al. 2018) its role and contribution are still a matter of debate.

In the computational neuroscience literature, the thalamus has been reported as a promising candidate for pattern classification analysis (Pergola et al. 2015). In addition, changes in the network topology of schizophrenic patients makes hubs ideal candidates for machine learning techniques as demonstrated by Cheng and colleagues (Cheng et al. 2015) with high accuracy when classifying patients and controls.

These are exciting outcomes since the declared goal of computational psychiatry is to provide physicians with tools that enable them to objectively identify patients where most approaches had been subjective up until that point. The hope is that the computational approach could be used to predict the likelihood of a previously unseen patient having schizophrenia. Identifying topological markers could lead to tools that enable to quantitatively determine the severity of common symptoms and even identify and measure the progression of the disease, as well as the effectiveness of treatment. Hence, it would be of scientific interest to find prognostic and diagnostic markers, not only based on empirical observations but also on outcomes obtained by a graph theoretical analysis. Our future research will focus on potential biomarkers and classification in schizophrenia.

Comparing the FD network and pure functional connectivity network based on pFC, we showed in the present study that the FD model is able to select links, of neurobiological meaning for schizophrenia, that otherwise would be neglected by the pFC analysis. Hence, the FD model stresses relevant features of the whole brain connectivity by adding structural as well as topological information to pFC. Therefore, we suggest that the FD model could be considered as an additional effective model to describe functional neural networks.

We believe that increasing the precision of the anatomical data in the model will reveal even further connectivity differences. In addition, it might contribute to a better understanding about the relationship between structural modularity and functional modularity of the brain, an open and challenging problem. Lastly, it is important to stress that the FD model is not only limited to the modality of fMRI. It can be also applied to PET (Positron Emission Tomography), EEG (electro-encephalography) and MEG (magneto-encephalography) data.

## Acknowledgements

CGF would like to thank Dr. Leonie Ascone Michelis for fruitful discussion about the role of the thalamo-cortical network in schizophrenia.

SK has been funded by two grants from the German Science Foundation (DFG KU 3322/1-1, SFB 936/C7), the European Union (ERC-2016-StG-Self-Control-677804) and a Fellowship from the Jacobs Foundation (JRF 2016-2018). CGF was funded by the German Science Foundation (SFB 936/C7). LK has been funded by the Evangelisches Studienwerk Villigst.

Authors contribution
Data acquisition: LK, JB, LS, PG, MF, NK, CM, SK, JG
Data analysis: CGF, PF, AP, SK, PD
Manuscript writing: CGF, PF, AP, JG, SK, PD

